# Mutual interplay of actin meshwork and stress fibers in cellular adaptive response: Insights from percolation dynamics

**DOI:** 10.1101/2023.12.24.573252

**Authors:** Yuika Ueda, Daiki Matsunaga, Shinji Deguchi

## Abstract

Cells dynamically remodel their internal structures by modulating the arrangement of actin filaments (AFs). In this process, individual AFs exhibit stochastic behavior without knowing macroscopic higher-order structures they are meant to create or disintegrate. Cellular adaptation to environmental cues is accompanied with this type of self-assembly and disassembly, but the mechanism allowing for the stochastic process-driven remodeling of the cell structure remains incompletely understood. Here we employ percolation theory to explore how AFs interacting only with neighboring ones without recognizing the overall configuration can nonetheless construct stress fibers (SFs) at particular locations. To achieve this, we determine the binding and unbinding probabilities of AFs undergoing cellular tensional homeostasis, a fundamental property maintaining intracellular tension. We showed that the duration required for the assembly of SFs is shortened by the amount of preexisting actin meshwork, while the disassembly occurs independently of the presence of actin meshwork. This asymmetry between the assembly and disassembly, consistently observed in actual cells, is explained by considering the nature of intracellular tension transmission. Thus, our percolation analysis provides insights into the role of coexisting higher-order actin structures in their flexible responses during cellular adaptation.

## 1. Introduction

One of the salient features of the cell structure is its adaptive capability to remodel the architecture according to the roles required in a given environment. In eliciting this capability, individual actin filaments (AFs), serving as the primary structural component, exhibit stochastic behavior without “knowledge” of the macroscopic structure they are meant to create (Yamashiro et al., 2018; Miller et al., 2022; Saito et al., 2022). AFs instead merely interact with neighboring elements, culminating in the collective construction of a cell structure comprised of specific actin-based structures (Chrzanowska-Wodnicka and Burridge, 1996; Hotulainen and Lappalainen, 2006). It also implies the potential of cells to update the internal structure by modulating the probability of the interaction, that is binding and unbinding, as well as the quantity or characteristics of the components. Currently, the mechanism behind this flexible creation of the cell structure utilizing the stochastic behavior of individual components remains incompletely understood.

The above perspective, focusing on the cellular hierarchy comprised of structural components interacting with each other, evokes a concept in physics known as percolation theory (Broadbent and Hammersley, 1957). Percolation theory describes how stochastic phenomena at a microscopic level yield an order at a macroscopic level by examining the interconnection among nodes within the system. Numerous probabilistic processes have been explored in diverse fields of science with the framework of percolation, which include invasion percolation, oriented percolation, and bond percolation (Alvarado et al., 2013; Langley et al., 2018). The percolation theory has drawn the interest of physicists because of its simple setup as well as non-trivial phenomena it exhibits in bridging different hierarchies. On the other hand, while some aspects of the cell structure have been scrutinized through the approach of percolation theory (Forgacs, 1995; Silveira et al., 2009; Pritchard et al., 2014; Ni et al., 2022), its adaptive nature driven by environmental cues remains largely unknown.

Among actin structures in mesenchymal non-muscle cells such as fibroblasts, actin meshwork serves as a fundamental element providing essential structural support to the cytoplasm and mediating biochemical signals accordingly (Deguchi et al., 2005; Svitkina, 2020; Vignaud et al., 2021). Meanwhile, stress fibers (SFs) are expressed particularly on a stiff substrate while physically connecting separate focal adhesions (FAs) at their termini. SFs are comprised of bundles of AFs, which are cross-linked by non-muscle myosin II to exert contractile tension onto the FAs (Okamoto et al., 2019). In cells residing on less stiff substrates, directional SFs appear in the cytoplasm containing more disordered actin meshwork. Thus, there is an exchange of the constituent AFs between the coexisting structures, actin meshwork and SFs, allowing the cells to adapt to the given environment (Ueda et al., 2022). The coexistence of the two distinct actin structures has been discussed from a percolation perspective by attributing the relative abundance to a single probability that characterizes the interaction between them, but not yet thoroughly analyzed (Forgacs, 1995; Silveira et al., 2009). In particular, significant ambiguity persists regarding the biological role of the coexistence of actin-based structures in the cellular adaptive response to environmental stimuli.

Here we perform Monte Carlo simulations based on a percolation model to investigate how the stochastic behavior of AFs can lead to the distinction between directional SFs and random actin meshwork while also accommodating with environmental changes. The model incorporates the binding and unbinding probabilities of AFs considering the influence of cellular tensional homeostasis, a fundamental property of proliferative non-muscle cells (Hahn and Schwartz, 2009; Kaunas and Deguchi, 2011; Ueda and Deguchi, 2023). Tensional homeostasis allows for keeping a constant level of intracellular tension through turnover, and hence it is also referred to as mechanical homeostasis. With this function, mechanotransduction-mediated biochemical signals activated upon the tension are also maintained constant, circumventing chronic activation of pro-inflammatory signaling and hence facilitating the ability of cells to adapt to applied mechanical stress. Our percolation model encompasses AFs and their higher-order structures, actin meshwork and SFs, without explicitly considering numerous other components that actually exist in real cells such as non-muscle myosin II and filopodia/lamellipodia (Liu et al., 2022). Regardless of this simplicity inherent to percolation theory focusing on the most essential aspects, our primary goal here is not to mimic such actual complex phenomena but rather to extract the role of the coexistence of the higher-order structures in cellular adaptation. We also consider a minimal analytical model to delineate the distinction between the responses to elevation and reduction in homeostatic tension observed in the percolation analysis.

## 2. Model analysis

### 2.1 Distribution of AFs

The actual cells are comprised of a variety of molecules and are thus so complicated; but, to focus on the indispensable factor associated with intracellular force transmission, we explicitly consider only the involvement of AFs. Specifically, individual AFs are modeled to be a line segment of a length *L* and are distributed randomly in a two-dimensional *x* − *y* plane that represents a cell with a square shape. The position of one end of AFs (*x*_1_, y_1_) is given randomly within the cell plane at each time step, and the other end (*x*_2_, *y*_2_) is expressed by

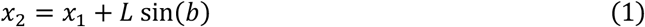

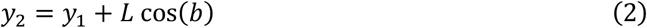

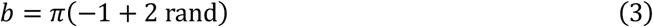

where *b* is the radian angle of the line segment, and rand is provided with a value between 0 and 1 according to a uniform random number generator.

### 2.2 Association and dissociation of AFs

Overlapping AFs stochastically bind with each other in the cell to potentially form a bundle of AFs defined as SFs. The AFs that constitute SFs undergo continuous turnover, namely molecular exchange with the surrounding actin meshwork, and thus the association and dissociation with the existing SFs occur simultaneously (Saito et al., 2022). To mimic this feature, the association and dissociation probabilities are defined as follows:

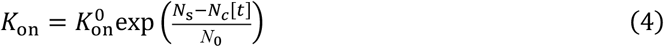

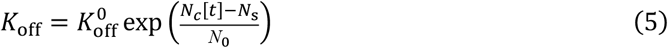

where *N*_s_ and *N*_*c*_[*t*] represents the number of AFs that is determined by the steady state tension in a SF undergoing tensional homeostasis and the actual number of AFs that constitute a SF at time *t*, respectively; N_0_ is a normalizing coefficient set to be 100; and, 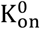 and 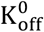are the association and dissociation probabilities at the initial state and set to be 0.35 and 0.20 (Wang et al., 2003; Kovács et al., 2007), respectively. These equations suggest that *K*_on_ is exponentially increased if the actual number of AFs constituting a SF is lower than that under steady state tensional homeostasis, while *K*_off_ is rather increased if the actual number is, on the contrary, larger than that at tensional homeostasis. Note that there is no deterministic situation in intracellular activities so that both of the probabilities will never be zero. The rationale behind this formula is that SFs are thickened in response to increased tension (Chrzanowska-Wodnicka and Burridge, 1996; Hirata et al., 2007). This is allowed by the catch bond properties, in which the lifetime of nonmuscle myosin on AFs is extended upon tension to eventually increase their number associated with SFs (Kovács et al., 2007; Okamoto et al., 2020). Thus, *N*_*c*_[*t*] is supposed to increase over time with tension development. Meanwhile, N_s_ refers to a specific number of AFs that is more intrinsic to the cell state. Specifically, the phosphorylation of myosin regulatory light chain (MRLC) dictates the potential allowing for the nonmuscle myosin–AF interaction, and the phosphorylation level varies depending on the intracellular and extracellular milieu (Matsui et al., 2019; Huang et al., 2021). For example, cells on softer substrate exhibit lower levels of MRLC phosphorylation, and it ends up in lower levels of tension and hence thinner SFs or lower numbers of associated AFs. Given this intrinsic property of cells, N_s_ is given as a parameter interpreted as a function of the MRLC phosphorylation reflecting the intrinsic cell state.

### 2.3 Analysis of SF, FAs, and actin meshwork

So far we only mentioned that SFs are bundles of AFs, and FAs are located at the termini of SFs; for the present model, we more specifically define that SFs are a state of AFs that align within a specific angle (−π/3 ≤ θ ≤ *π*⁄3) and collectively link two separate FAs; FAs are represented by two different points, each of which is located at the center of the facing edges of the square cell plane; and, θ is the angle measured from the line connecting the separate FAs. The rationale for the directional alignment of AFs is that the actual SFs are comprised of bundled AFs bearing tension, distinct from the highly entangled actin meshwork. Thus, consistent direction of the constituents should be considered to describe SFs given that having large numbers of AFs can end up making a link between the separate FAs even with no particular alignment. We assumed that n = 10 AFs are linked at its one end to each one of the FAs all the time, while the direction of the individual AFs can change randomly over time. These specific AFs are considered to constitute a part of the FAs and are thus not counted as being within the population of AFs. The actin meshwork is defined here to be a state of AFs that collectively link two facing edges of the cell plane that do not contain the FAs. Thus, unlike the case of SFs, the direction of individual AFs is not considered to specify actin meshwork given its randomly distributed nature.

### 2.4 Monte-Carlo simulation

The distribution of AFs is first determined based on Eqs. (1)–(3), and then *K*_on_ and *K*_off_ are determined based on Eqs. (4)–(5) by counting the number of AFs involved in the SF, i.e., *N*_−_[*t* = 0]. The state of AFs at the following time steps is determined by the Monte-Carlo method with the probabilities of *K*_on_ and *K*_off_. Specifically, AFs stay at their same positions with *K*_on_, while they change their positions again randomly with *K*_off_. The subsequent evolution of the position of AFs is analyzed by arbitrary numbers of steps regarded to be time *t*. At each time step *t*, the presence and absence of the AF-mediated link between the FAs yields 1 and 0, respectively. As the time evolution is dependent on the initial position of AFs and the stochastic event, the same Monte-Carlo simulation was repeated by 10^4^ to obtain the Percolation Probability (PP) that represents the mean of the probability to make a bridge between the FAs, or the probability to form a SF, at each time *t*. Likewise, PP for forming actin meshwork, which links facing edges of the cell plane as described above, is obtained.

### 2.5 Determination of the number of AFs

To determine *N*_*c*_[*t*], i.e., the number of AFs involved in the SF at time *t*, a line segment intersection detection algorithm using vector cross product is employed. Specifically, let one end of an AF, as a line segment, be *P*_1_(*x*_1_, *y*_1_) and the other end be *P*_2_(*x*_2_, *y*_2_); likewise, let *P*_3_(*x*_3_, *y*_3_) and *P*_4_(*x*_4_, *y*_4_) describe the two ends of another AF. The condition that the AFs cross with each other is described by using the vector cross product as

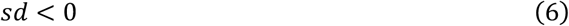

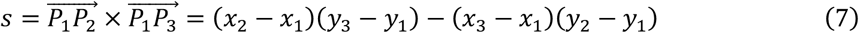

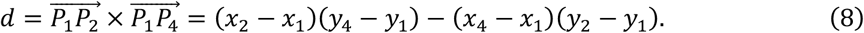

The presence of intersection is thus analyzed for all the combinations of individual AFs to count the total number associated with the FAs. AFs orienting within an angle of −π/3 ≤ θ ≤ *π*⁄3 are picked up in this process as described above to finally obtain *N*_*c*_[*t*] as the number constituting a straight SF.

## 3 Results

### 3.1 The effect of cellular tension generation potential on percolation

First, the effect of changing N_s_ on *N*_*c*_[*t*]was investigated. As already described, N_s_ represents a specific number of AFs that reflects the MRLC phosphorylation level or the inherent potential of cells to be able to generate a tension, while *N*_*c*_[*t*]is the actual number of AFs that constitute SFs. The length of individual AFs and each cell edge were set to be a constant 0.2 and 1.0, respectively. The number of individual AFs was set to be 250. All AFs are involved only in actin meshwork at *t* = 0 because they are initially distributed randomly in the cell (Fig. 1). AFs tend to remain in between the separate FAs to form a SF particular with a high value of N_s_. We analyzed how the number of individual AFs forming a SF changes over time as a function of N_s._ At a low N_s_ of 80 where the potential to genera a tension is relatively low, the majority of AFs remains as a part of actin meshwork without building SFs. Here, note that the sum of the different populations for actin meshwork and SFs remains constant at 250 over time. As N_s_ is augmented, e.g., to 150, the extent of the association of AFs with SFs is increased accordingly. Finally, upon a further augmentation of N_s_ to, e.g., 200, the dominant population of AFs reverses from that for actin meshwork to that for SFs. The number of AFs reaches a steady state depending on the value of N_s_, suggesting that actin meshwork and SFs stably coexist.

**Figure 1.**
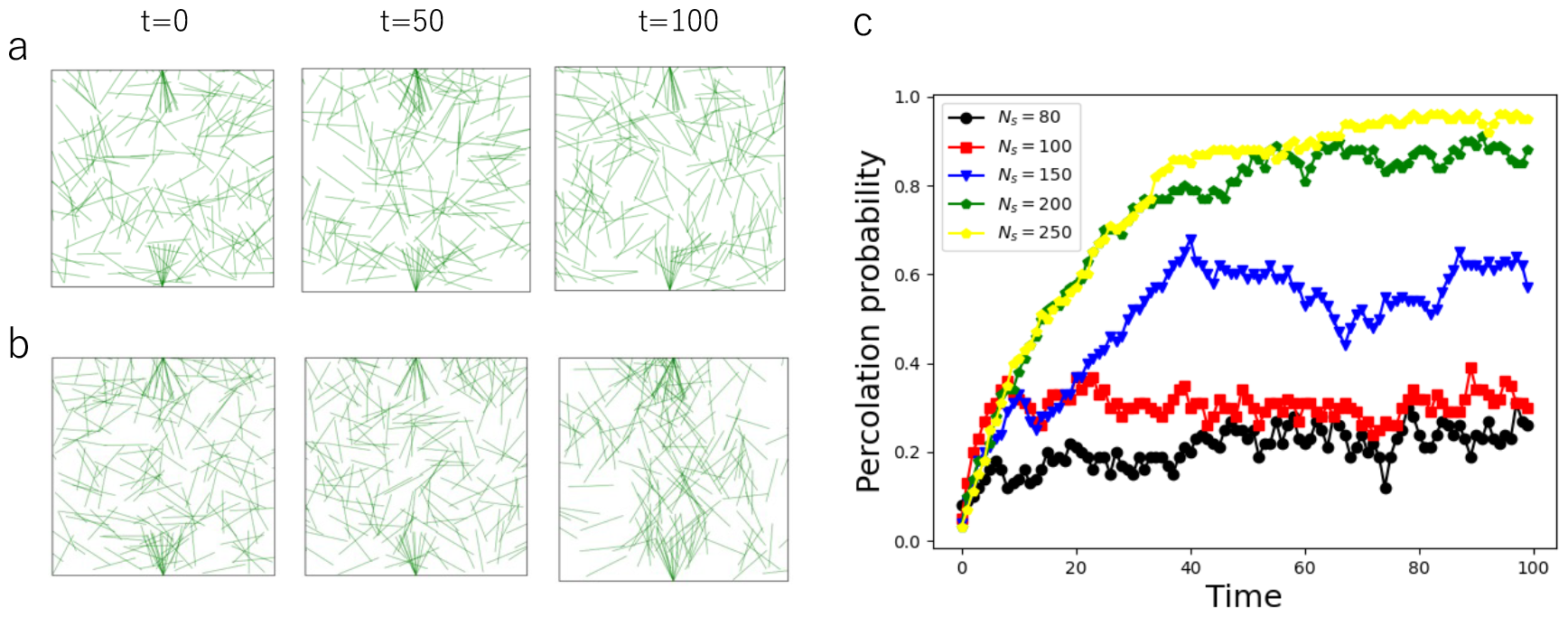
Percolation analysis. (a, b) Schematic model of the cell with two dimensional square lattices. AFs are randomly distributed in the cell. We evaluate the formation of SFs by calculating the connection between the two FAs located at the top and bottom edges of the cell. Distribution states of AFs at t = 0, t = 50, and t = 100for N_s_ = 80(a) and N_s_ = 200(b). (c) Time evolution of the percolation probability for N_s_ = 80(black),N_s_ = 100(red),N_s_ = 150(blue), N_s_ = 200(green), and N_s_ = 250(yellow). With a large N_s_, the probability for forming SFs converges to a large value.

The above data are actually a representative of a Monte-Carlo-based stochastic process. To analyze how the Percolation Probability (PP) changes over time, where PP measures the probability of whether AFs form a SF or not, we repeated the simulation 10^4^ times for each of N_s_ = 80, 100, 150, 200, and 250. The average behavior shows that PP gradually increases over time because the lifetime of AFs on a SF is extended upon binding due to the *K*_on_-mediated catch bond properties to allow the SF to be stabilized. The SF finally reaches a steady state where the association and dissociation are approximately balanced while subject to constant turnover. A greater value of PP was obtained at the steady state with a higher value of N_s_, which raises *K*_on_ and lowers *K*_off_ to stabilize the existing SF.

### 3.2 The effect of AF length distribution on percolation

The need for having specific AF lengths is often focused on at the level of single intracellular structures such as primary cilia and muscle sarcomeres (Littlefield et al., 2021; Macarelli et al., 2023). For example, sarcomeres are optimized for efficient contraction by expressing nebulin to keep a fixed AF length (Labeit et al., 2011). Meanwhile, only little discussion has been made on the necessary length of individual AFs at the level of whole cells. However, given that filamin is distributed over the cytoplasm to convert local tension into biochemical signals related to tensional homeostasis (DuFort et al., 2011), the regulation for AF lengths may exist at the whole cell level as well. We previously investigated based on statistical mechanics and thermodynamics the physical requirement for allowing cells to maintain tensional homeostasis (Ueda et al., 2022). Specifically, we assumed that AFs as a tension-bearing component must have as diverse lengths as possible and occupy as much space as possible in the cytoplasm in accordance with the requirement for maximum entropy production. The theoretical consequence was that AFs need to have more short subpopulations than long ones, and in fact the length distributions obtained were in qualitative agreement with those that have been observed in experiments. Other potential biological advantages for possessing such specific AF length distributions were not fully discussed in these studies. With the present numerical approach to analyze the intracellular percolation, we are thus in a position to tackle this open question of how the distribution relates to the function of cells.

The effect of providing a variety of the length of individual AFs, namely *L*_0_, on PP was thus investigated, while keeping the same total length of all AFs, or in other words the same number of actin molecules. First, we determined the length of individual AFs according to the exponential distribution,

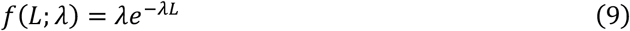

where *λ* is a parameter that characterizes the distribution. The rationale for employing the exponential distribution is that previous studies including ours have indicated that the length of AFs inside of living cells has an exponential distribution-like probability function (Matsui et al., 2011; Ueda et al., 2022). Here, we gave *λ* a value of 8, 6, or 4 to have a distribution inclined to short length regions to a highest (Fig. 2a), middle (Fig. 2b), or weakest degree (Fig. 2c), respectively. Each cell edge was set to be 1.0 as in the above analysis. The time course of PP shows that the most uniform length distribution (*λ* = 4) and the one most inclined to shorter length regions (*λ* = 8) allow SFs and actin meshwork to be relatively easily formed, respectively, while the middle one (*λ* = 6) exhibits a tendency in between them (Fig. 2c, d). Thus, the length distribution affects the preferred choice of cross-linking style in the cytoplasm, in which actin meshwork requires tension transmission all throughout the cytoplasm so that short AF populations suitably contribute to the formation, whereas SFs bear a tension in particular directions so that the presence of long AFs would be efficient and enough to do that.

**Figure 2.**
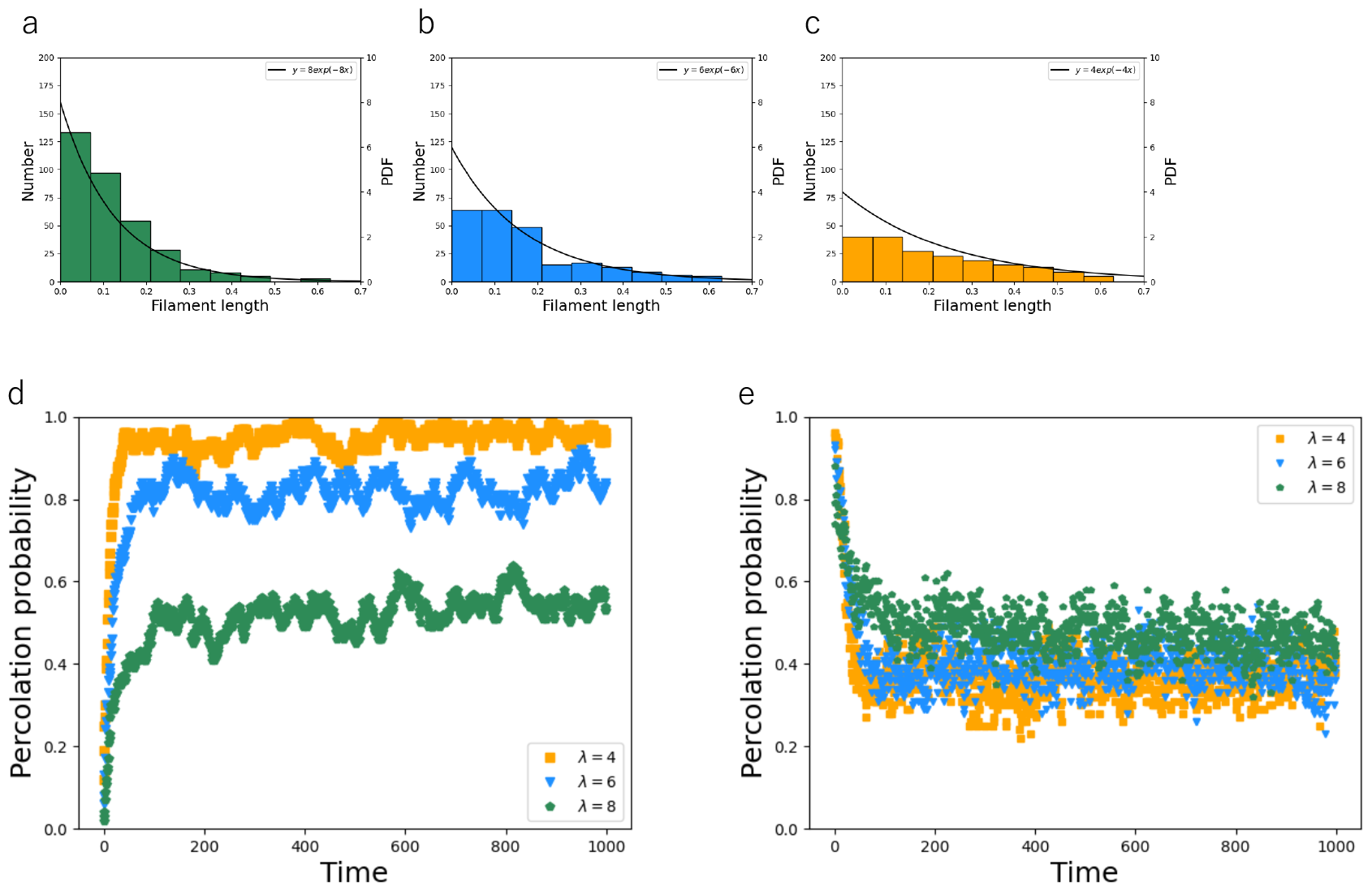
The effect of AF length distribution on percolation probability. (a, b, c) Probability density function and number of AFs at different length distributions. y = 8e^−8*x*^ (a),y = 6e^−6*x*^ (b), and y = 4e^−4*x*^ (c). (d) Time evolution of the percolation probability of SFs.(e) Time evolution of the percolation probability of actin meshwork.

### 3.3 Adaptive response to change in cellular tension generation potential

Next, the effect of changing the value of N_s_ in the process of percolation development was investigated, which aims at analyzing the adaptive response of the cell model to environmental change. As N_s_ reflects the cellular capability of generating a tension, which varies in accordance with the surrounding milieu, the temporal change in N_s_ simulates a process in which cells are exposed to pharmacological, signaling, or biophysical stimulation; e.g., cells increase and decrease the capability of generating a tension on stiff and soft substrates as well as with increased and decreased Rho-kinase activity as the kinase for MRLC, respectively (Matsui et al., 2019). For this simulation, the length of individual AFs and each cell edge were set to be 0.2 and 1.0, respectively, and the number of individual AFs was set to be 250 as in the above analysis.

First, we analyzed the effect of decreasing the value of N_s_ from 150 or 200 to 80 at *t* = 60 where the system approximately reaches the steady state after the development of percolation initiated at *t* = 0(Fig. 3a). The states of N_s_ = 150 and 200 represent the cell where actin meshwork and SFs coexist and SFs exist predominantly over actin meshwork, respectively, and then the response to a stepwise decrease to 80 where actin meshwork should be dominant was analyzed. The results indicate that PP sharply decreases upon the chage in N_s_ in both cases. Next, we increased the value of N_s_, on the contrary, from 80 or 150 to 200 in a stepwise manner at *t* = 60(Fig. 3b). Interestingly, unlike the above case that PP decreases at apparently the same rate upon the change in N_s_, the current case with the increase in N_s_ from 80 to 200 takes longer time compared to that from 150 to 200. To evaluate the difference in the rate of the response, all the PP values were collected during an adaptive period of between *t* = 60 and 150 for each condition and were analyzed with student *t*-test. The analysis shows that there is no significant difference in the PP values during the adaptive period between the change of N_s_ from 200 to 80 and that from 150 to 80. Meanwhile, the PP values were significantly higher for the change of N_s_ from 150 to 200 compared to that from 80 to 200. Thus, the extent of the coexistence of actin meshwork and SFs does not affect the rate of percolation collapse, while it does affect the rate of percolation formation, in which advanced coexistence (i.e., initial N_s_ = 150) is advantageous for rapid adaptation to complete percolation than less developed conditions (i.e., initial N_s_ = 80).

**Figure 3.**
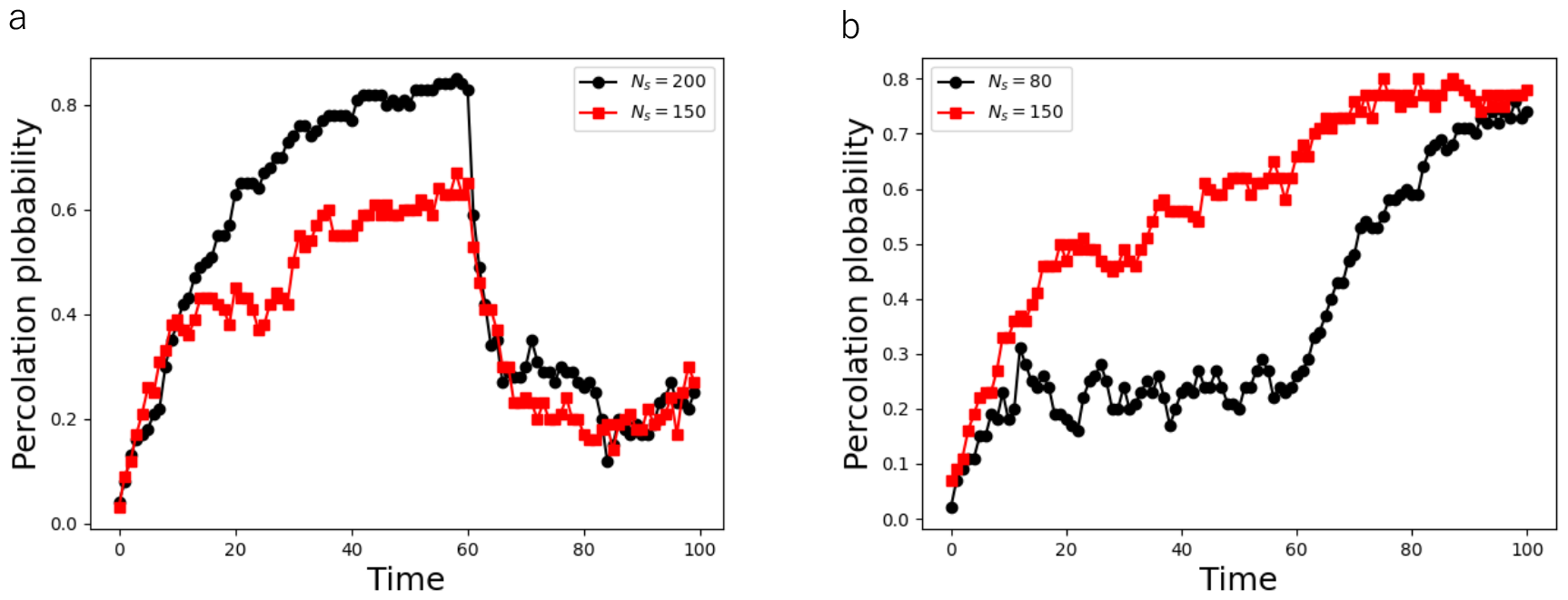
Adaptive response of the percolation nature. (a) Response to change in N_s_ for N_s_ = 200(black) and N_s_ = 150(red). (b) Response to change in N_s_ for N_s_ = 80(black) and N_s_ = 150(red).

## 4 Discussion

What is intriguing and essential in general percolation analyses is the consideration of a hierarchical system comprising at least two layers, in which the individual elements of the lower layer interact with neighboring elements at a local level without recognizing the overarching structure, and then only modulating the stochastic rate of these interactions at the microscopic level can end up in an emergence of a new structure or a phase-transition of the system at the macroscopic level (Ueda et al., 2023). These hierarchical systems can be observed in individual adherent cells, in which tension plays a pivotal role in mechanosensitive regulation in the actin meshwork-based cytoplasm to form SFs at suitable locations (Deguchi et al., 2011). Cells thus flexibly reconfigure their internal structures in accordance with the surrounding milieu, exhibiting an inherent percolation nature in their adaptive behavior.

More specifically, directional SFs are created from and maintained within the random actin meshwork (Vignaud et al., 2021), acting as a physical connector between separate FAs in a process reminiscent of general percolation. Our percolation analysis thus simulated the stochastic interaction among AFs as a microscopic component and the potential emergence of SFs as a macroscopic structure spanning the entire cell. The probability of association between the tension-bearing element AFs is modulated by N_s_ that represents the cellular capacity to generate homeostatic tension, which is basically determined in actual cells by each cell type and environment, thus serving as a global parameter. There is another factor that locally modulates the probability, namely *N*_*c*_[*t*], which denotes the number of AFs associated with an existing SF. In other words, individual AFs interact only with coincidentally overlapped and bound AFs via tension transmission without recognizing the overarching cell structure, but nonetheless can construct SFs at particular locations.

Thus, the states of microscopic elements and their collective entity are coordinately interrelated within the hierarchical cell system, and a change in one layer impacts the others by leveraging the inherent nature or percolation to flexibly cope with continuously generating intracellular/extracellular perturbations. This percolation-based perspective is essential in comprehending the environment-dependent self-assembly of living systems composed of multiple interacting components. This adaptive nature seems crucial for proliferative cells that are supposed to remodel their internal structure to alleviate the stress inevitably provided from time-varying intracellular and extracellular milieu (Hahn and Schwartz, 2009; Kaunas and Deguchi, 2011). In contrast, this percolation-based architectural maintenance differs from more organized structures found in non-proliferative cells such as muscle cells, which, as already described above, express nebulin to stabilize the length of AFs and are specialized to contract only unidirectionally so that the muscle cells, unlike non-muscle cells, are not required to address indistinct mechanical stresses exerted from various directions (Labeit et al., 2011; Ueda et al., 2022).

Our percolation analysis showed that a high and low level of N_s_ leads to a different cell state where SFs and actin meshwork are more preferably expressed, respectively. These results agree well with experimental observations given that N_s_ is associated with the level of MRLC phosphorylation (Matsui et al., 2019). N_s_ in reality reflects more general capability of the cell to generate a homeostatic tension so that this parameter is actually affected by various environmental cues. For example, cells more preferably express actin meshwork and SFs on a softer and stiffer substrate and at a smaller and larger adhesive area, respectively (Prager-Khoutorsky et al., 2011), and this selective architectural style can also be interpreted to correspond in our percolation analysis to a case with a low and high value of N_s_.

We further found that the coexistence of SFs with a larger amount of actin meshwork delays the time to reach a steady state following elevation in N_s_, while it causes no time delay in relation to reduction in N_s_ regardless of the actin meshwork amount. In other words, the assembly of SFs is influenced by the amount of preexisting actin meshwork, while the disassembly occurs rapidly independent of the presence of actin meshwork. This asymmetry between the assembly and disassembly is reminiscent of typical metabolic processes, in which anabolism often involves a relatively prolonged period of time, while catabolism is promptly induced. Consistently, SFs in actual cells can be disassembled more rapidly than their assembly (Matsui et al., 2010; Matsui et al., 2011; Huang et al., 2021; Saito et al., 2021). This tendency can be explained by focusing in our model on that AFs associated with SFs are the elements responsible for transmission of tension or specifically the information of K_on_. Meanwhile, AFs belonging to actin meshwork merely continue to move randomly. Thus, SFs must first be created, if cells are initially comprised only of actin meshwork, for the transmission of mechanical information and then be increased in number to finally reach a steady state. From a biological perspective, the importance of actin meshwork can thus be interpreted that the flexible response of non-muscle cells to a dynamically changing environment is partly allowed by the presence of actin meshwork unlike muscle cells possessing distinct actin bundles that are supposed to persistently perform a single defined role to sustain a tension in the defined axial direction.

## Availability of data and materials

All data generated or analyzed during this study are included in this published article and its supplementary information files.

## Authors’ contributions

Y.U. and S.D. conceived research; Y.U. designed and conducted research with feedback from S.D.; D.M. provided technical support; Y.U. and S.D. wrote the manuscript. All authors read and approved the final manuscript.

## References

Alvarado, J., et al. Molecular motors robustly drive active gels to a critically connected state. Nature Physics 9 (2013): 591–597.

Broadbent, S.R., Hammersley, J.M. Percolation processes: I. Crystals and mazes. Mathematical proceedings of the Cambridge philosophical society. Vol. 53. No. 3. Cambridge University Press, 1957.

Chrzanowska-Wodnicka, M., Burridge, K., J Cell Biol (1996) 133 (6): 1403–1415.

Deguchi, S., Ohashi, T., Sato, M., Intracellular stress transmission through actin stress fiber network in adherent vascular cells. MCB Molecular & Cellular Biomechanics 2, 4, 205-216, 2005.

Deguchi, S., Matsui, T.S., Iio, K., The position and size of individual focal adhesions are determined by intracellular stress-dependent positive regulation, Cytoskeleton, 68, 639–651, 2011.

DuFort, C.C., Paszek, M.J., Weaver, V.M. Balancing forces: architectural control of mechanotransduction. Nature Reviews Molecular Cell Biology 12, 308–319, 2011.

Forgacs, G., On the possible role of cytoskeletal filamentous networks in intracellular signaling: an approach based on percolation. J Cell Sci (1995) 108 (6): 2131–2143.

Hahn, C., Schwartz, M.A. 2009. Mechanotransduction in vascular physiology and atherogenesis. Nat Rev Mol Cell Biol. 10(1), 53–62.

Hirata, H., Tatsumi, H., Sokabe, M., Dynamics of actin filaments during tension-dependent formation of actin bundles. Biochimica et Biophysica Acta (BBA) - General Subjects 1770, 8, 2007, 1115–1127.

Hotulainen, P., Lappalainen, P., J Cell Biol (2006) 173 (3): 383–394.

Huang, W., Matsui, T.S., Saito, T., Kuragano, M., Takahashi, M., Kawahara, T., Sato, M., Deguchi, S. Mechanosensitive myosin II but not cofilin primarily contributes to cyclic cell stretch-induced selective disassembly of actin stress fibers. Am J Physiol Cell Physiol 320, C1153–C1163, 2021.

Kaunas, R., Deguchi, S. 2011. Multiple roles for myosin II in tensional homeostasis under mechanical loading. Cell Mol Bioeng 4 (2):182–191.

Kovács, M., Thirumurugan, K., Knight, P.J., Sellers, J.R., Load-dependent mechanism of nonmuscle myosin 2. Proc Natl Acad Sci U S A 104, 24, 9994-9999, 2007.

Labeit, S., Ottenheijm, C.A.C., Granzier, H. 2011. Nebulin, a major player in muscle health and disease. FASEB J 25(3):822–829.

Langley, D.P., et al. Percolation in networks of 1-dimensional objects: comparison between Monte Carlo simulations and experimental observations. Nanoscale Horizons 3 (2018): 545–550.

Littlefield, R., Almenar-Queralt, A., Fowler, V.M., Actin dynamics at pointed ends regulates thin filament length in striated muscle. Nature Cell Biology 3, 544–551, 2001.

Liu, S., Matsui, T.S., Kang, N., Deguchi, S., Analysis of senescence-responsive stress fiber proteome reveals reorganization of stress fibers mediated by elongation factor eEF2 in HFF-1 cells. Molecular Biology of the Cell 33, 1, ar10, 2022.

Macarelli, V., Leventea, E., Merkle, F.T., Regulation of the length of neuronal primary cilia and its potential effects on signalling. Trends in Cell Biology 33, 11, 979-990, 2023.

Matsui, T.S., Ito, K., Kaunas, R., Sato, M., Deguchi, S., Actin stress fibers are at a tipping point between conventional shortening and rapid disassembly at physiological levels of MgATP, Biochemical and Biophysical Research Communications, 395, 301–306, 2010.

Matsui, T.S., Kaunas, R., Kanzaki, M., Sato, M., Deguchi, S. Non-muscle myosin II induces disassembly of actin stress fibres independently of myosin light chain dephosphorylation. Interface Focus 1, 5, 754-766, 2011.

Matsui, T.S., Deguchi, S. Spatially selective myosin regulatory light chain regulation is absent in dedifferentiated vascular smooth muscle cells but is partially induced by fibronectin and Klf4. Am J Physiol Cell Physiol 316, C509–C521, 2019.

Miller, C.M., et al., Nature Communications volume 13, 4749, 2022.

Ni, Q., et al., A tug of war between filament treadmilling and myosin induced contractility generates actin rings. eLife 11, 2022.

Okamoto, T., Matsui, T.S., Ohishi, T., Deguchi, S., Helical structure of actin stress fibers and its possible contribution to inducing their direction-selective disassembly upon cell shortening. Biomechanics and Modeling in Mechanobiology 19, 543–555, 2020.

Prager-Khoutorsky, M., Lichtenstein, A., Krishnan, R., Rajendran, K., Mayo, A., Kam, Z., Geiger, B., Bershadsky, A.D., Fibroblast polarization is a matrix-rigidity-dependent process controlled by focal adhesion mechanosensing. Nat Cell Biol 13,12, 1457–1465, 2011.

Pritchard, R.H., Huang, Y.Y.S., Terentjev, E.M., Mechanics of biological networks: from the cell cytoskeleton to connective tissue. Soft Matter, 2014, 10, 1864–1884.

Saito, T., Huang, W., Matsui, T.S., Kuragano, M., Takahashi, M., Deguchi, S., What factors determine the number of nonmuscle myosin II in the sarcomeric unit of stress fibers? Biomechanics and Modeling in Mechanobiology 20, 155–166, 2021.

Saito, T., Matsunaga, D., Deguchi, S., Analysis of chemomechanical behavior of stress fibers by continuum mechanics-based FRAP. Biophysical Journal 121, 2921–2930, 2022.

Saito, T., Matsunaga, D., Deguchi, S., Long-term molecular turnover of actin stress fibers revealed by advection-reaction analysis in fluorescence recovery after photobleaching. PLoS ONE 17, 11, e0276909, 2022.

Silveira, P.S.P., Alencar, A.M., Majumdar, A., Lemos, M., Fredberg, J.J., Suki, B., Percolation in a network with long-range connections: Implications for cytoskeletal structure and function. Physica A: Statistical Mechanics and its Applications 388, 8, 2009, 1521–1526.

Svitkina, T.M., Actin cell cortex: structure and molecular organization. Trends in Cell Biology 30, 7, 556-565, 2020.

Ueda, Y., Matsunaga, D., Deguchi, S., A statistical mechanics model for determining the length distribution of actin filaments under cellular tensional homeostasis. Scientific Reports 12, 14466, 2022.

Ueda, Y., Deguchi, S., Emergence of multiple set-points of cellular homeostatic tension. Journal of Biomechanics 151, 111543, 2023.

Vignaud, T., Copos, C., Leterrier, C., Toro-Nahuelpan, M., Tseng, Q., Mahamid, J., Blanchoin, L., Mogilner, A., Théry, M., Kurzawa, L., Stress fibres are embedded in a contractile cortical network. Nature Materials 20, 410–420, 2021.

Wang, F., Kovács, M., Hu, A., Limouze, J., Harvey, E.V., Sellers, J.R., Kinetic mechanism of nonmuscle myosin IIB functional adaptations for tension generation and maintenance. Journal of Biological Chemistry 278 (2003): 27439–27448.

Yamashiro, S., et al., Molecular Biology of the Cell Vol. 29, No. 16, 2018, 1917–2035.

